# Genome-wide CRISPR screens identify DNA repair and R-loop suppression as regulators of the cellular sensitivity to environmentally relevant Bisphenol A exposure

**DOI:** 10.64898/2026.04.13.718249

**Authors:** Anastasia Hale, Alexandra Nusawardhana, Joshua Straka, Claudia M. Nicolae, George-Lucian Moldovan

## Abstract

Bisphenol A (BPA) is a prevalent chemical used in the production of plastics. While adverse effects on the reproductive system have been documented, more recent studies also associated BPA exposure with carcinogenesis as well as genomic instability. However, these studies were generally performed using BPA concentrations much higher than those observed in the serum or urine of the general population, making their relevance unclear. To address this, we report here an unbiased genetic study to identify mechanisms responding to environmentally relevant BPA exposure. We performed genome-wide CRISPR knockout screens in HeLa and RPE1 cells upon continuous exposure to 0.5uM BPA, a concentration similar to the mean BPA concentration found in the urine of plastics manufacturing workers, for 19 days. We found genome stability genes among the top common hits between the two cell lines, suggesting that BPA causes DNA damage at this environmentally relevant exposure dose. We validated the DNA repair gene RAD51C and the RNA helicase DDX21 as genes required for BPA resistance. Moreover, we show that BPA exposure increases the formation of R-loops which are resolved by DDX21. Our study suggests that BPA exposure at environmentally relevant doses can cause DNA damage, highlighting the relevance of BPA for carcinogenesis.

## Introduction

Bisphenol A (BPA) is a high production volume chemical involved in the manufacturing of polycarbonate plastics. It is found in products such as plastic food containers, water bottles, lining of metal food cans, water supply pipes, and dental sealant. Exposure to BPA is pervasive. In 2003-2004, the National Health and Nutrition Examination Survey (NHANES III) conducted by the Centers for Disease Control and Prevention (CDC) detected BPA in 93% of 2517 urine samples from individuals six years and older [1]. As of 2021, the median creatinine-adjusted urinary BPA concentration in adults was estimated to be 1.76 μg/g (95% PI: 0.79-2.73), and the pooled estimate for serum BPA was 1.75 μg/l (95% PI: 0-10.58) [2]. These values are even greater in those employed in the plastic manufacturing industry, with urinary BPA concentrations ranging from 0.78–18900 µg/g [3]. Moreover, no current OSHA standards exist for the plastics industry, making protections for these workers from the biochemical effects of BPA virtually nonexistent.

Exposure to BPA has been associated with a wide variety of adverse health effects such as reproductive abnormalities, placental dysfunction, precocious puberty, and neurological impairment [4,5]. More recently, BPA has been associated with carcinogenesis. For example, a study in 2020 showed that Ontario women employed in rubber manufacturing plants exhibited higher rates of lung and breast cancer [6]. However, the status of BPA as carcinogen remains controversial.

The mechanisms of carcinogenesis previously explored focused partially on the activation of estrogen signaling, particularly in the context of altered mammary gland development and proliferation. BPA is an endocrine disrupting chemical which has been shown to activate the estrogen receptor (ER). Previously explored mechanisms of BPA-mediated carcinogenesis in the context of ER signaling include enhanced activation of the Src/Raf/Erk pathway and subsequent chromosome mis-segregation, as well as increased expression of HOXB9, a homeobox-containing gene that plays an important role in mammary gland development [7]. Activation of the ER has also been shown to promote genomic instability. For example, ER-mediated transcription has been shown to induce the formation of cell-cycle dependent double stranded DNA breaks (DSBs) [8]. Moreover, estrogen (E2) mediated ER activation has been shown to promote the formation of co-transcriptional DNA-RNA hybrids (or R-loops), which have been linked to DNA damage [9]. BPA has also been shown to promote R-loop formation in ER-positive cells [10]. Whether BPA-mediated formation of R-loops also occurs in ER-negative cells is unclear.

The mechanism of action of many environmental carcinogens involves formation of DNA adducts, which cause mutations during DNA replication. Currently, it is under debate if BPA causes DNA damage [7]. Previous studies have shown that BPA exposure can cause formation of DNA adducts. However, these studies have been performed using BPA concentrations significantly higher than those relevant for the exposure of the general population to BPA. For example, several studies have identified the formation of BPA adducts in rat liver and murine mammary tissue, albeit at concentrations of 200mg/kg, approximately 100-fold greater than those identified in human serum [11,12]. Moreover, cell culture studies showed that BPA treatment causes single stranded DNA (ssDNA) gaps and DSBs [13,14]. However, these studies were also performed with high concentrations of BPA (100-200μM) for short durations. A recent study [13] explored the mutational signature associated with BPA exposure in 293T cells and found increased point mutagenesis centered on guanine residues, in line with previous studies showing that BPA preferentially forms guanosine adducts [15,16]. In addition, the authors also found an increase in chromosomal translocations, suggesting the formation of DSBs upon BPA exposure. Importantly, the authors also found that the mutational signatures observed in these cell-based studies bear similarities with several cancer-associated mutational signatures, suggesting that BPA exposure may be associated with carcinogenesis through DNA adduct formation. However, this study was performed be exposing cells to a high dose of BPA (100μM, about 20,000-fold higher than the mean serum concentration in the general public) for a short time (24h).

Considering the debate regarding the relevance of BPA exposure to DNA damage-induced carcinogenesis, we designed a genetic study to unbiasedly interrogate the biological processes regulating the cellular sensitivity to a physiologically-relevant concentration of BPA, and investigate if BPA exposure at environmentally relevant levels may cause DNA damage and activate DNA repair pathways. We performed genome-wide CRISPR knockout screens in HeLa and RPE1 cells upon continuous exposure to 0.5uM BPA, a concentration similar to that found in the urine of plastics manufacturing workers [3], for 19 days. We found DNA repair, cell cycle, sister chromatid cohesion, and chromosome instability genes among the top common hits between the two cell lines, suggesting that BPA causes DNA damage at this environmentally relevant exposure. We validate the DNA repair gene RAD51C and the RNA helicase DDX21 as genes required for BPA resistance, show that BPA exposure potentially promotes R-loop formation even in ER-negative cells, and that these R-loops are possibly resolved by DDX21. Our study suggests that BPA exposure at environmentally relevant doses can cause DNA damage, highlighting the relevance of BPA for carcinogenesis.

## Methods

### Cell culture and protein techniques

HeLa and RPE1-p53KO cells were grown in Dulbecco’s modified Eagle’s media (DMEM). Gene depletion by sgRNA was performed using commercially available CRISPR/Cas9 KO plasmids: RAD51C (Santa Cruz Biotechnology sc-403060), DDX21 (Santa Cruz Biotechnology sc-405770-KO-2), ATG9A (Santa Cruz Biotechnology sc-408011), CPSF2 (Santa Cruz Biotechnology sc-407012).

### Western Blot

Denatured whole cell extracts were prepared by boiling cells in 100mM Tris, 4% SDS, 0.5M β-mercaptoethanol. Protein samples were loaded and separated on an SDS-PAGE gel and transferred to a PVDF nitrocellulose membrane (ThermoFisher Scientific). After blocking with 5% milk in TBST, the membrane was incubated with a solution containing the primary antibody in 5% milk at 4°C overnight. The membrane was then incubated with a HRP-conjugated secondary antibody in 5% milk for an hour. Antibodies used for Western blot, at 1:500 dilution, were:

RAD51C: Santa Cruz Biotechnology sc-56214;

DDX21: Cell Signaling Technology 59278;

ATG9A: Cell Signaling Technology 13509;

CPSF2: Invitrogen PA5-31004;

ER-α: Abcam ab108398;

GAPDH: Santa Cruz Biotechnology sc-47724;

Vinculin: Santa Cruz Biotechnology sc-73614.

### CRISPR screens

The Brunello Human CRISPR knockout pooled lentiviral library (Addgene 73179) was used for CRISPR knockout screens [17]. This library encompasses 76,411 gRNAs that target 19,114 genes. 55 million cells from each cell lines were infected with the Brunello CRISPR library at a multiplicity of infection (MOI) of 0.4 to achieve 250-fold coverage and selected for 4 days with 0.6μg/mL puromycin. Twenty million library-infected were passaged cells (to maintain 250-fold coverage) for 14 days in the presence of DMSO or 0.5μM BPA and then collected. Genomic DNA was isolated using the DNeasy Blood and Tissue Kit (Qiagen 69504) and employed for PCR using Illumina adapters to identify the gRNA representation in each sample. 10μg of gDNA was used in each PCR reaction along with 20μl 5X HiFi Reaction Buffer, 4μl of P5 primer, 4μl of P7 primer, 3μl of Radiant HiFi Ultra Polymerase (Stellar Scientific), and water. The P5 and P7 primers were determined using the user guide provided with the CRISPR libraries (https://media.addgene.org/cms/filer_public/61/16/611619f4-0926-4a07-b5c7-e286a8ecf7f5/broadgpp-sequencing-protocol.pdf). The PCR cycled as follows: 98°C for 2min before cycling, then 98°C for 10sec, 60°C for 15sec, and 72°C for 45sec, for 30 cycles, and finally 72°C for 5min. After PCR purification, the final product was Sanger sequenced to confirm that the guide region is present, followed by qPCR to determine the exact amount of PCR product present. The purified PCR product was then sequenced with Illumina HiSeq 2500 single read for 50 cycles, targeting 10 million reads. Next, the sequencing results were analyzed bioinformatically using the MAGeCK algorithm, which takes into consideration raw gRNA read counts to test if individual guides vary significantly between the conditions [18]. The MAGeCK software and instructions on running it were obtained from https://sourceforge.net/p/mageck/wiki/libraries/. The ranking lists of genes obtained for each screen comparison are presented in Supplementary Tables S1, S2 and S3. Finally, analyses of the Gene Ontology and biological processes enriched among the top hits was performed using DAVID [19,20].

### Functional assays

For cellular survival assays, 500,000 cells were passaged every 3-4 days in 10cm plates in the presence of either a DMSO vehicle control or 0.5μM of BPA for 21 days. Total number of cells per plate were calculated at each passage, and the projected cell count was determined. Immunofluorescence was performed as previously described [21] using the following primary antibodies: γH2AX (MilliporeSigma JBW301); 53BP1 (Fortis A300-272A); DDX21 (Cell Signaling Technology 59278). Slides were imaged on a confocal microscope (Leica SP5) and analyzed using ImageJ 1.53a software.

### Proximity ligation-based assays

For PLA assays, cells were treated with DMSO or 0.5μM BPA for 48 hours and seeded into 8-chamber slides. 0.5μM BPA was added for another 24 hours, after which cells were permeabilized with 0.5% Triton for 10min at 4°C, washed with PBS, fixed at room temperature with 3.7% paraformaldehyde in PBS for 10min, washed again in PBS and then blocked in Naveni blocking solution (Navinci Diagnostics NT.1.100.01) for 1hr at 37°C, and incubated overnight at 4°C with primary antibodies. Antibodies used were: S9.6 (Kerafast ENH001), dsDNA (Novus Biologicals NBP3-07302). Samples were then subjected to a proximity ligation reaction using the NaveniFlex Cell MR kit (Navinci Diagnostics NC.MR.100) according to the manufacturer’s instructions. Slides were imaged using a confocal microscope (Leica SP5) and images were analyzed using ImageJ 1.53a software.

### Fluoresence image acquisition and analysis

Fluorescence images were acquired using a confocal microscope (Leica SP5 STED) with the HC PL APO ×63/1.40 oil objective utilizing the Type F Immersion liquid (11944399, Leica Microsystems). The 405-nm laser was used for DAPI (cell nuclei staining) while White Light Laser (WLL) was used for Alexa Fluor Plus 488 (53BP1 staining), Alexa Fluor Plus 568 (γH2AX and DDX21 staining), and Texas Red 594 (SIRF and PLA foci) excitations. HyD detectors were used for a subsequent fluorescence detection. Leica Application Suit X (LAS X) software (Leica Microsystems) was used for optimization of fluorescence detection, signal yields, and spectral unmixing. All images were taken without overexposure at the identical laser intensity, gain, and exposure parameters. The images were saved as 2048 pixels × 2048 pixels, 8-bit multi-channel Leica Image Files (.lif) and exported for quantification to 8-bit TIFF format files using LAS X software. γH2AX, 53BP1, DDX21, PLA, and SIRF foci were quantified using Image J 1.53a software. Nuclei were outlined via the ‘analyze particles’ function, and a uniform prominence was set using the ‘Find Maxima’ function to detect foci for each antibody. Representative images for figures were prepared using Image J and exported as 8-bit TIFF format files. Brightness was linearly adjusted for a given fluorochrome uniformly across entire images and figures to enhance visibility while ensuring the data are accurately represented.

### Statistical analyses

For immunofluorescence and PLA assays the t-test (two-tailed, unpaired) was used. For cellular viability assays, Wilcoxon matched-pairs signed rank test (two-tailed) was used. Statistical analyses were performed using GraphPad Prism 10. Statistical significance is indicated for each graph (ns = not significant, for p>0.05; * for p≤0.05; ** for p≤0.01; *** for p≤0.001, **** for p≤0.0001). The random probabilities of identical genes within the top hits with MAGeCK score lower than 0.01 were calculated by multiplying the individual probabilities of each set: [(number of genes in set 1/total number of genes in the library)*(number of genes in set 2/total number of genes in the library)].

## Results

### CRISPR screens to identify genetic determinants of the cellular response to low-dose BPA exposure

To identify the genetic mechanisms responding to BPA exposure, we performed a set of unbiased CRISPR screens in cervical cancer HeLa cells and non-cancer RPE1 epithelial cells. We employed the Brunello CRISPR-knockout lentiviral library [17], which targets 19,114 human genes and averages 4 guide RNAs (gRNAs) for each gene, resulting in a total of 76,441 unique gRNAs. After selection, maintaining 250x fold library coverage (equivalent to 20 million cells) at all times, we treated library-infected cells with 0.5µM BPA or DMSO vehicle control for 19 days, splitting cells every 3 days with BPA added freshly every time (Figure 1A). The BPA concentration used is similar to the median BPA concentration found in the urine and serum of plastics manufacturing workers.

**Figure 1.**
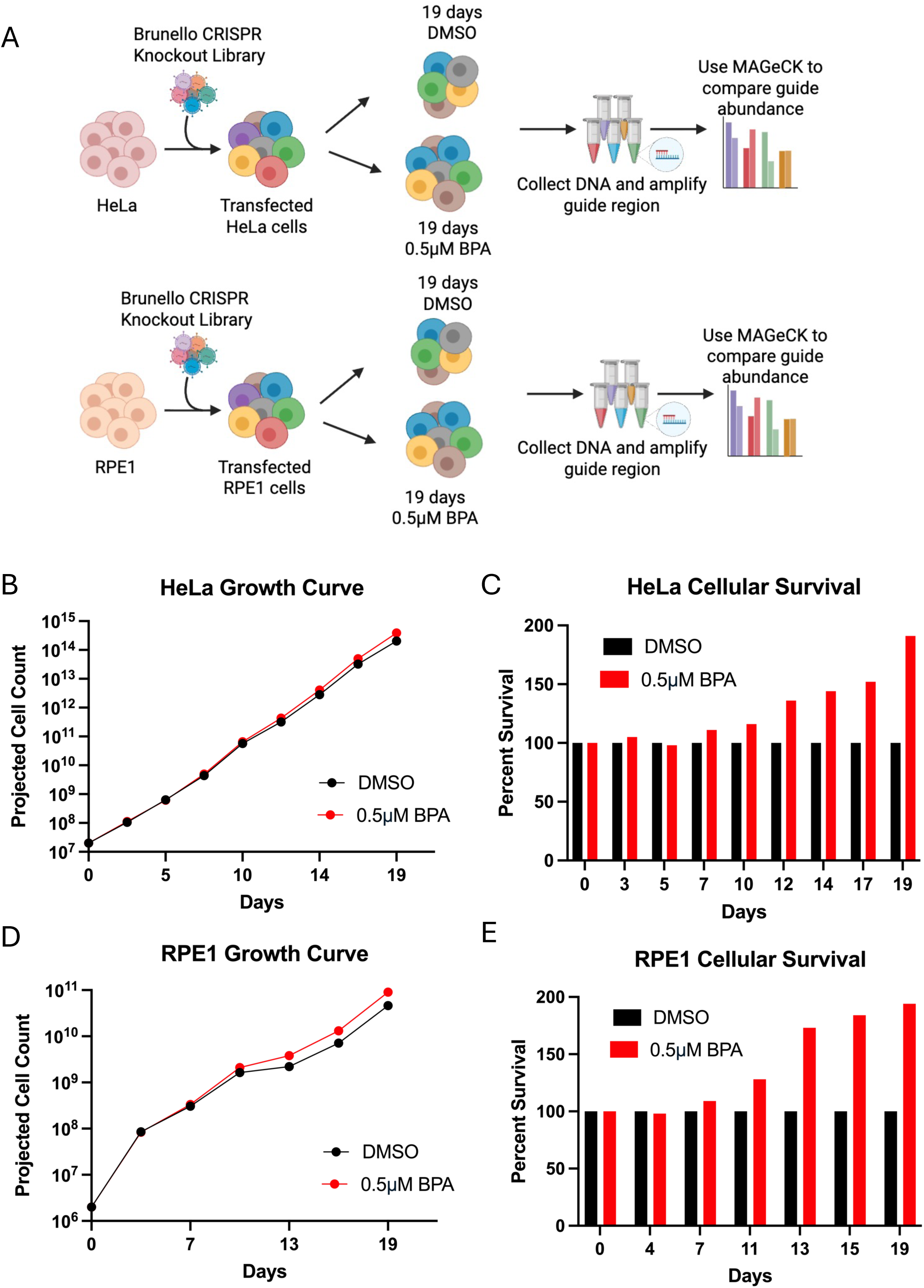
Genome-wide CRISPR knockout screens for BPA sensitivity. (A) Overview of the CRISPR knockout screens to identify genes that are specifically required for cells exposed to BPA. HeLa and RPE1 cells were incubated with and without 0.5 μM BPA for 19 days before being collected and analyzed via MAGeCK. Created in BioRender. Moldovan, G. (2026) https://BioRender.com/t1z1vw4. (**B-E**) Analyses of cellular proliferation throughout the screen. (**B, D**) Graphs showing projected cell count from day 0 to day 19 for Brunello library-infected HeLa (**B**) and RPE (**D**) cells treated with bisphenol A (BPA) and DMSO treated control cells. Cells were counted at each split and projected cell count was calculated to assess the impact of BPA exposure on cell proliferation. (**C, E**) Graphs showing percent survival (based on projected cell count) from day 0 to day 19 for HeLa (**C**) and RPE (**E**) cells treated with bisphenol A (BPA) compared to the DMSO treated control at each time point.

We noted that continuous treatment with 0.5µM BPA of the library-infected cells slightly increased the proliferation of both HeLa and RPE1 cells. This effect started after 10 days of treatment and became more and more pronounced with each passage after that (Figure 1B-E).

At the end of the BPA treatment, cells were collected, and genomic DNA was extracted. PCR was used to amplify the gRNA region and the gRNA was identified by Illumina sequencing. Bioinformatic analyses using the MAGeCK algorithm [42] were used to generate ranking lists of genes which were lost in BPA-treated cells compared to untreated cells (Supplementary Tables S1, S2). This represents genes which, when inactivated, result in increased cell death in BPA-treated cells compared to untreated cells.

We then analyzed these lists in order to assess the overlap between the screen results in the two cell lines. There were 64 common genes within the top hits with MAGeCK score lower than 0.01 (667 genes for the HeLa screen and 799 genes for the RPE1 screen) (Figure 2A). This is greater than the random probability of common genes within two datasets of these sizes, which is 28 (Figure 2B), suggesting common responses to BPA across the two cell lines.

**Figure 2.**
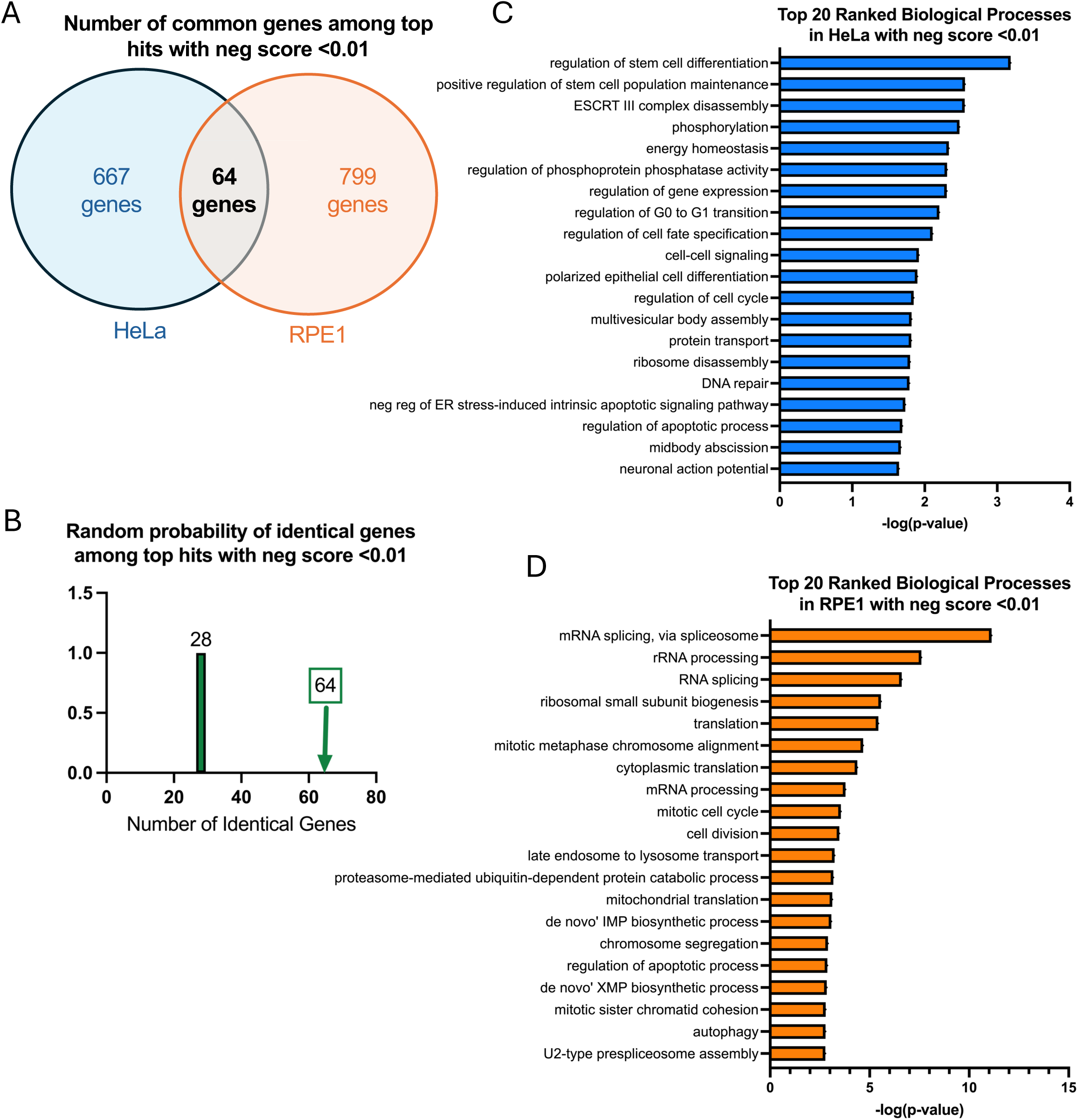
Analyses of the top hits from genome-wide CRISPR knockout screens for BPA sensitivity. (**A**) Diagram showing the overlap of identical genes within the top hits with MAGeCK score lower than 0.01 which cause sensitivity to BPA in HeLa and RPE1 cells. (**B**) The number of common genes within the top hits with MAGeCK score lower than 0.01 which cause sensitivity to BPA in HeLa and RPE1 cells (namely 64) is higher than the random probability of identical hits, which is 28. (**C, D**) Biological pathway analyses using Gene Ontology of the top hits with MAGeCK score lower than 0.01 which cause sensitivity to HeLa (C) and RPE1 (D) cells. GO_BP terms with negative logP greater than 1.64 (C) and 2.65 (D) are presented.

To gain insights into these responses, we performed biological pathway analyses of the top hits with MAGeCK score lower than 0.01, using Gene Ontology (Biological Processes). Surprisingly, this analysis yielded limited overlap between the biological processes identified in the two cell lines (Figure 2C,D). On careful examination, DNA-related processes appear to be the only ones in common (DNA repair, cell cycle, sister chromatid cohesion, chromosomal instability).

### Validation of top hits from the CRISPR screens for BPA sensitivity

We next sought to validate top hits from the CRISPR screens, to demonstrate their impact on cellular sensitivity upon BPA treatment. We focused on common hits which ranked high in both cell lines (Figure 3A-D; Supplementary Table S3). This included the top ranked genes CPSF2 (ranked 2 in the RPE1 screen and 69 in the HeLa screen) and ATG9A (ranked 6 in the RPE1 screen and 58 in the HeLa screen) (Figure 3E). We also investigated two genome stability genes found within the 64 common hits with MAGeCK score lower than 0.01, namely DDX21, an RNA helicase involved in R-loop resolution (ranked 97 in the HeLa screen and 719 in the RPE1 screen), and the DNA repair gene RAD51C (ranked 332 in the HeLa screen and 269 in the RPE1 screen).

**Figure 3.**
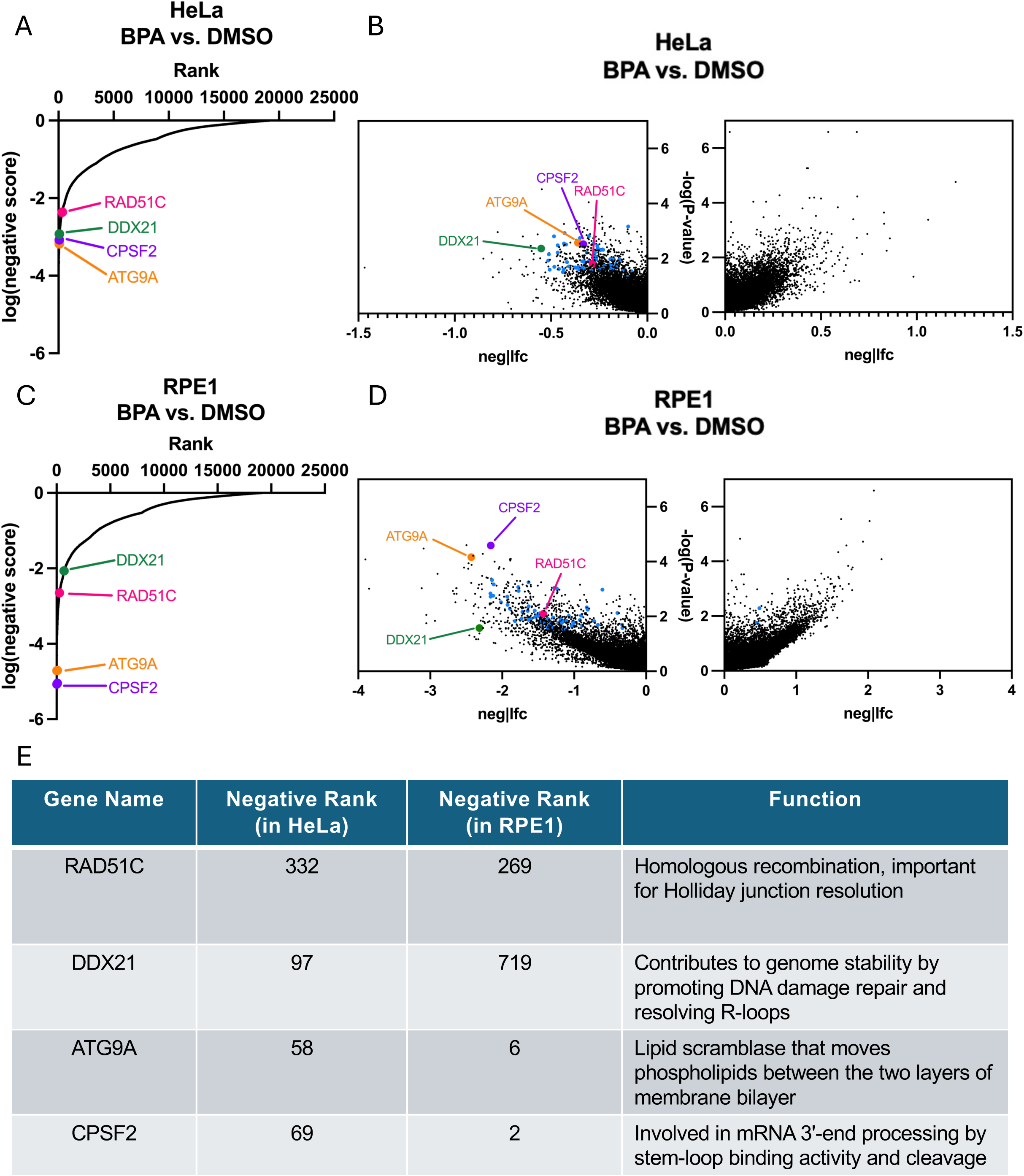
Selection of top hits from genome-wide CRISPR knockout screens for BPA sensitivity. (**A-D**) Plots of screens results. (**A, C**) Scatterplots showing the results of genome-wide CRISPR knockout screens to identify genes that are specifically required for sensitivity to BPA in HeLa (**A**) and RPE1 (**C**) cells. Four top hits chosen for validation are indicated. (**B, D**) Volcano plots showing the results of genome-wide CRISPR knockout screens to identify genes that are specifically required for sensitivity to BPA in HeLa (**B**) and RPE1 (**D**) cells. Genes targeted by the library are plotted by the −log10 of their respective negative and positive *P*-values and associated log2 fold change values with genes clustering to the upper left of the graph demonstrating a reduction in relative survival when lost in the BPA treated cells. Four top hits chosen for validation are indicated. (**E**) Table showing screens ranking, and the biological functions of the validated top hits in the BPA sensitivity screens.

Since we wanted to validate these hits using the same condition as those employed in the original screens, namely exposure to 0.5μM BPA for three weeks, we could not employ transient siRNA-mediated knockdown of the top hits. We thus instead used CRISPR-mediated gene inactivation for stable downregulation of the top hits. Using a lentiviral-mediated Cas9 and sgRNA delivery system, we were able to downregulate the expression of all 4 genes in RPE1 cells, as well as downregulate DDX21 and ATG9A depletion in HeLa cells. In most cases, the depletion was not complete, indicating that not all alleles were knocked out (Figure 4A). Despite our repeated efforts, we were not able to obtain better depletions of these genes. This perhaps reflect the fact that RAD51C, DDX21 and ATG9A are known to be embryonic lethal when knocked-out in mice [22,23] (The Jackson Laboratory entry MGI:1860494).

**Figure 4.**
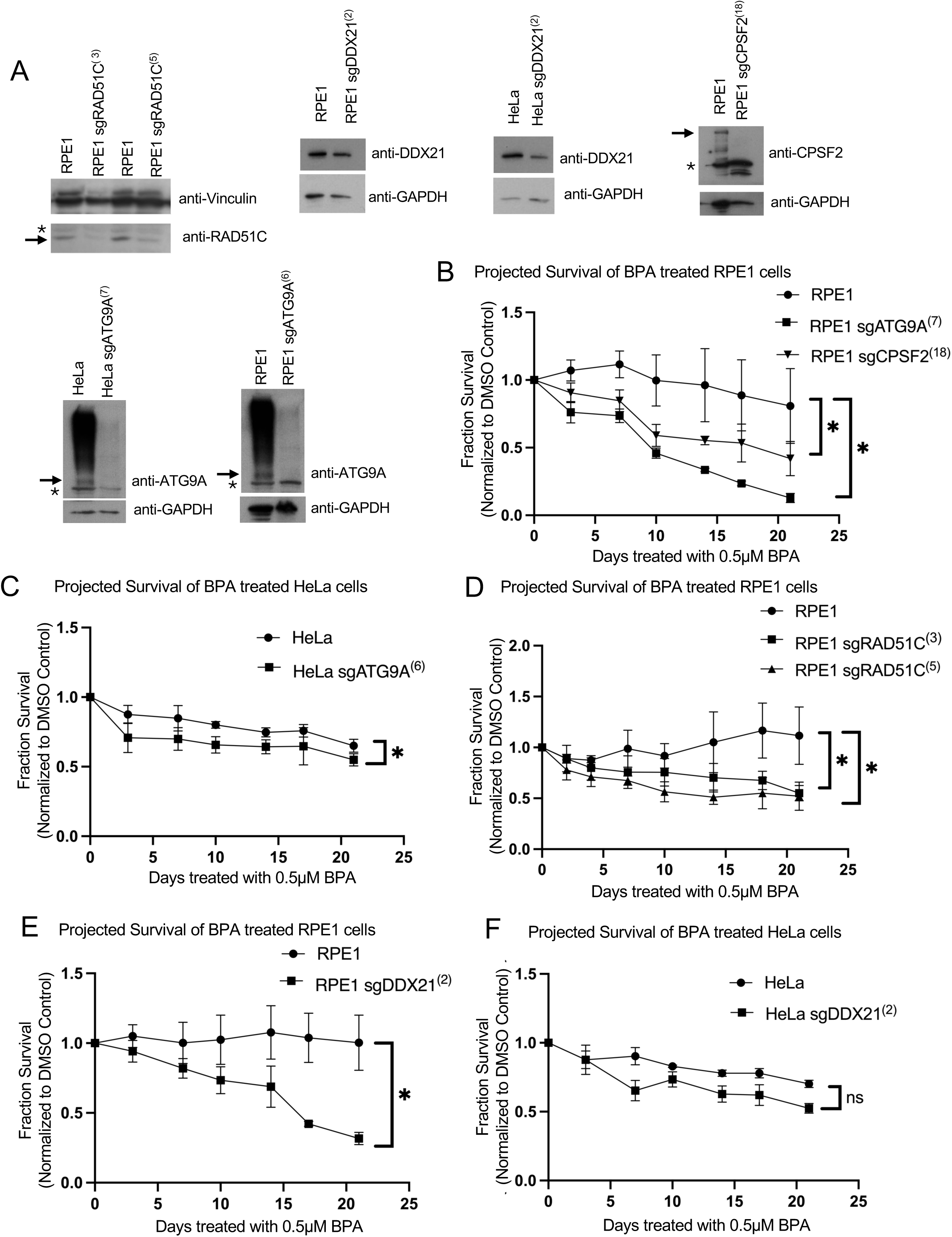
Validation of BPA sensitivity hits ATG9A, CPSF2, RAD51C, and DDX21. (**A**) Western blots showing sgRNA-mediated depletion of ATG9A, CPSF2, RAD51C, and DDX21. If multiple bands are present, arrows are used to indicate the respective proteins. Asterisks indicate cross-reactive bands. For the ATG9A blot, the high molecular weight smear likely corresponds to a heavily post-translationally modified form of the protein, as it disappears in the sgATG9A cells. (**B-F**) Cellular survival of CPSF2 and ATG9A-depleted RPE1 cells (**B**), ATG9A-depleted HeLa cells (**C**), RAD51C-depleted RPE1 cells (**D**), DDX21-depleted RPE1 cells (**E**), and DDX21-depleted HeLa cells (**F**) over 21 days of exposure to 0.5 μM BPA, normalized to DMSO treated controls. Averages of three independent replicates are plotted at each time point, and asterisks indicate statistical significance (Wilcoxon matched-pairs signed rank test, two-tailed).

We treated the sgRNA cell lines with 0.5μM BPA or DMSO vehicle control for 21 days, with fresh BPA added every 3-4 days when cells were passaged. Cell proliferation analyses indicated that RPE1 sgATG9A and sgCPSF2 cell lines showed increased BPA sensitivity compared to control cells (Figure 4B). Similarly, HeLa sgATG9A cells were more sensitive to BPA than wildtype cells (Figure 4C). Moreover, both RPE1 sgRAD51C clones we obtained showed increased BPA sensitivity compared to wildtype cells (Figure 4D). Finally, DDX21 sgRNA depletion in both HeLa and RPE1 resulted in BPA sensitivity under these conditions (Figure 4E,F). While statistically significant, for both sgDDX21 and sgATG9A the BPA sensitivity seemed to be more pronounced in RPE1 compared to HeLa cells, likely reflecting cell type-specific effects. Overall, these BPA sensitivity experiments validate our CRISPR screens and confirm that ATG9A, CPSF2, RAD51C and DDX21 are required for the cellular resistance to environmentally relevant BPA exposure.

### RAD51C suppresses BPA-induced DSBs

RAD51C is a RAD51 paralog involved in homologous recombination-mediated repair of DNA double stranded breaks [24]. Our results showing that loss of RAD51C causes increased cellular sensitivity upon exposure to 0.5μM BPA for 21 days suggest that, even at this low, environmentally-relevant concentration, BPA exposure can cause DNA damage and/or replication stress. To investigate this, we measured γH2AX foci, a marker of DSBs, using immunofluorescence experiments. Exposure to 0.5μM BPA for 3 days resulted in an increase in γH2AX foci (Figure 5A), confirming that exposure to low-dose BPA can cause DNA damage. We then investigated the impact of RAD51C depletion on γH2AX foci. Even without BPA treatment, RAD51C-depleted cells showed an increase in γH2AX foci (Figure 5A). This possibly reflects the role on RAD51C in repairing endogenous DNA damage. Surprisingly, unlike in wildtype cells, BPA treatment caused a reduction in γH2AX foci in both sgRAD51C clones we obtained (Figure 5A-D).

**Figure 5.**
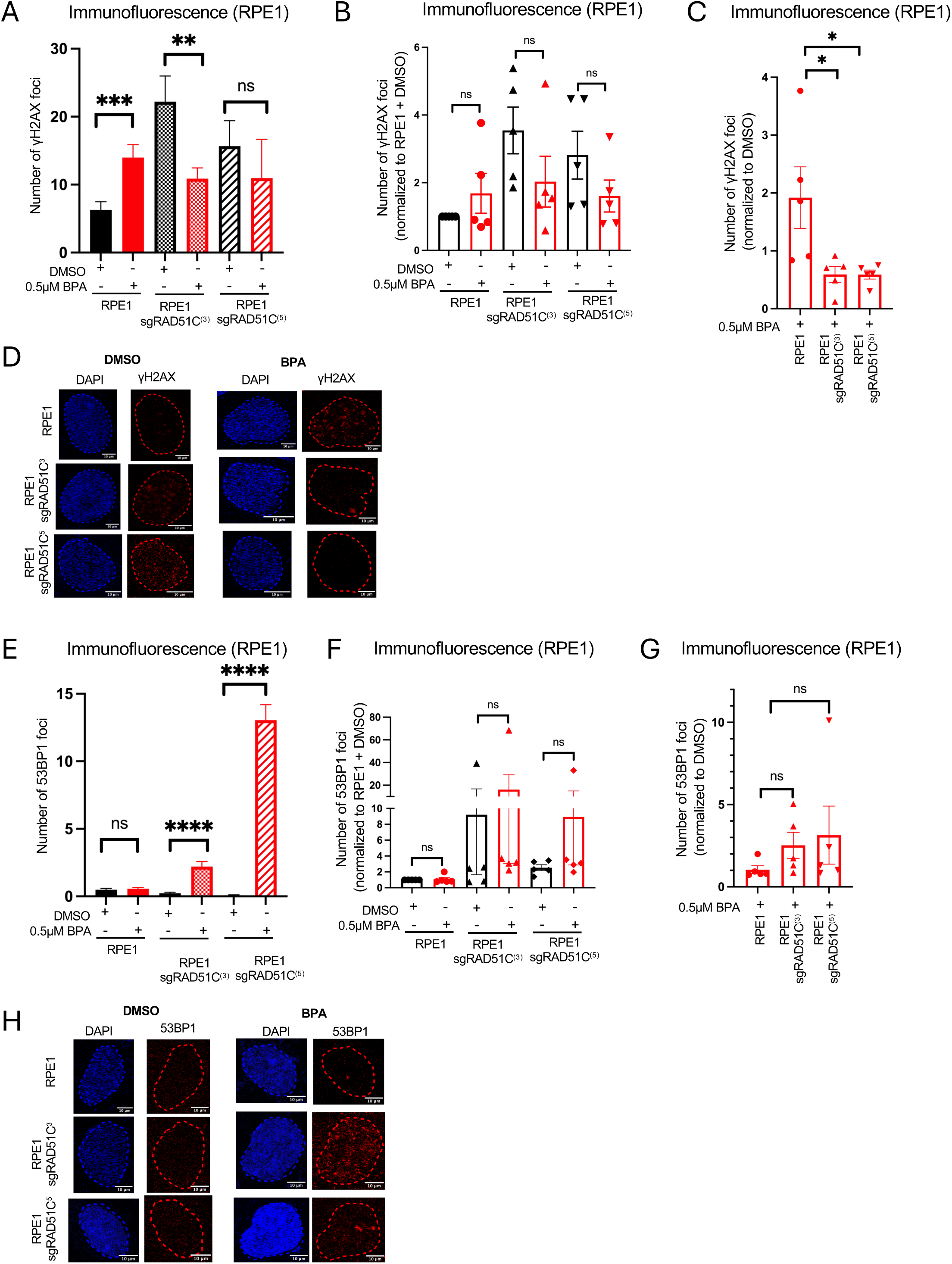
Environmentally relevant BPA exposure causes DSBs which are suppressed by RAD51C. (**A-D**) γH2AX immunofluorescence showing that treatment with 0.5 μM BPA for 72 hours increases γH2AX foci in in RPE1 cells but reduces γH2AX foci in RPE1 sgRAD51C cells as compared to DMSO treated controls. (**A**) Representative γH2AX immunofluorescence experiment. At least 67 cells were quantified for each condition. The mean value is represented on the graphs, and asterisks indicate statistical significance (t-test two-tailed, unpaired). (**B,C**) The means of 5 independent γH2AX immunofluorescence experiments, normalized to RPE1 + DMSO (**B**) and further normalized to DMSO control treatment for each cell line (**C**) are plotted (t-test two-tailed, unpaired). (**D**) Representative γH2AX immunofluorescence micrographs are shown. (**E-H**) 53BP1 immunofluorescence showing that treatment with 0.5 μM BPA for 72 hours increases 53BP1 foci formation in RPE1 sgRAD51C cells as compared to DMSO treated controls. (**E**) Representative γH2AX immunofluorescence experiment. At least 80 cells were quantified for each condition. The mean value is represented on the graphs, and asterisks indicate statistical significance (t-test two-tailed, unpaired). (**F,G**) The means of 5 independent 53BP1 immunofluorescence experiments, normalized to RPE1 + DMSO (**F**) and further normalized to DMSO control treatment for each cell line (**G**) are plotted (t-test two-tailed, unpaired). (**H**) Representative 53BP1 immunofluorescence micrographs are shown.

These findings may indicate that loss of RAD51C reduces the DNA damage induced by BPA. On the other hand, since γH2AX represents H2AX phosphorylated by DNA damage checkpoint kinases at DNA damage sites, these findings may instead indicate that, while DSBs may still form, the DNA damage response system is not correctly activated upon RAD51C depletion. Indeed, a previous study showed that RAD51C is required for the phosphorylation of CHK2 by ATM in response to DSBs [25]. To distinguish between these two possibilities, we measured 53BP1 foci, another marker of DSBs. 53BP1 binds to DSBs and initiates their repair by NHEJ, a potentially error-prone DSB repair pathway which can cause chromosomal translocations. We found that, in both sgRAD51C cell lines, treatment with 0.5μM BPA for 3 days caused an increase in 53BP1 foci (Figure 5E-G). While the difference is not statistically significant, the trend was consistently observed. Overall, these findings suggest that exposure to low-dose environmentally relevant BPA causes DNA damage, potentially double stranded DNA breaks. This damage can be repaired by homologous recombination. We speculate that, in RAD51C-depleted cells, DSB repair signaling and HR are compromised, resulting in increased DNA damage which is repaired through mutagenic 53BP1-mediated NHEJ, potentially resulting in chromosomal translocations.

### DDX21 suppresses the accumulation of BPA-induced R-loops in estrogen receptor-negative cells

DDX21 is an RNA helicase previously shown to suppress R-loops [26]. Since we found that DDX21 is required for cellular resistance to BPA, we first investigated if DDX21 is recruited to DNA upon BPA exposure. Using immunofluorescence experiments, we found that DDX21 forms nuclear foci, which are increased upon treatment with 0.5μM BPA for 3 days in both HeLa and RPE1 cells (Figure 6A,B). These foci appear to be peri-nucleolar, similar to the appearance of R-loops in previously-published immunofluorescence experiments using the S9.6 antibody recognizing DNA-RNA hybrids [27].

**Figure 6.**
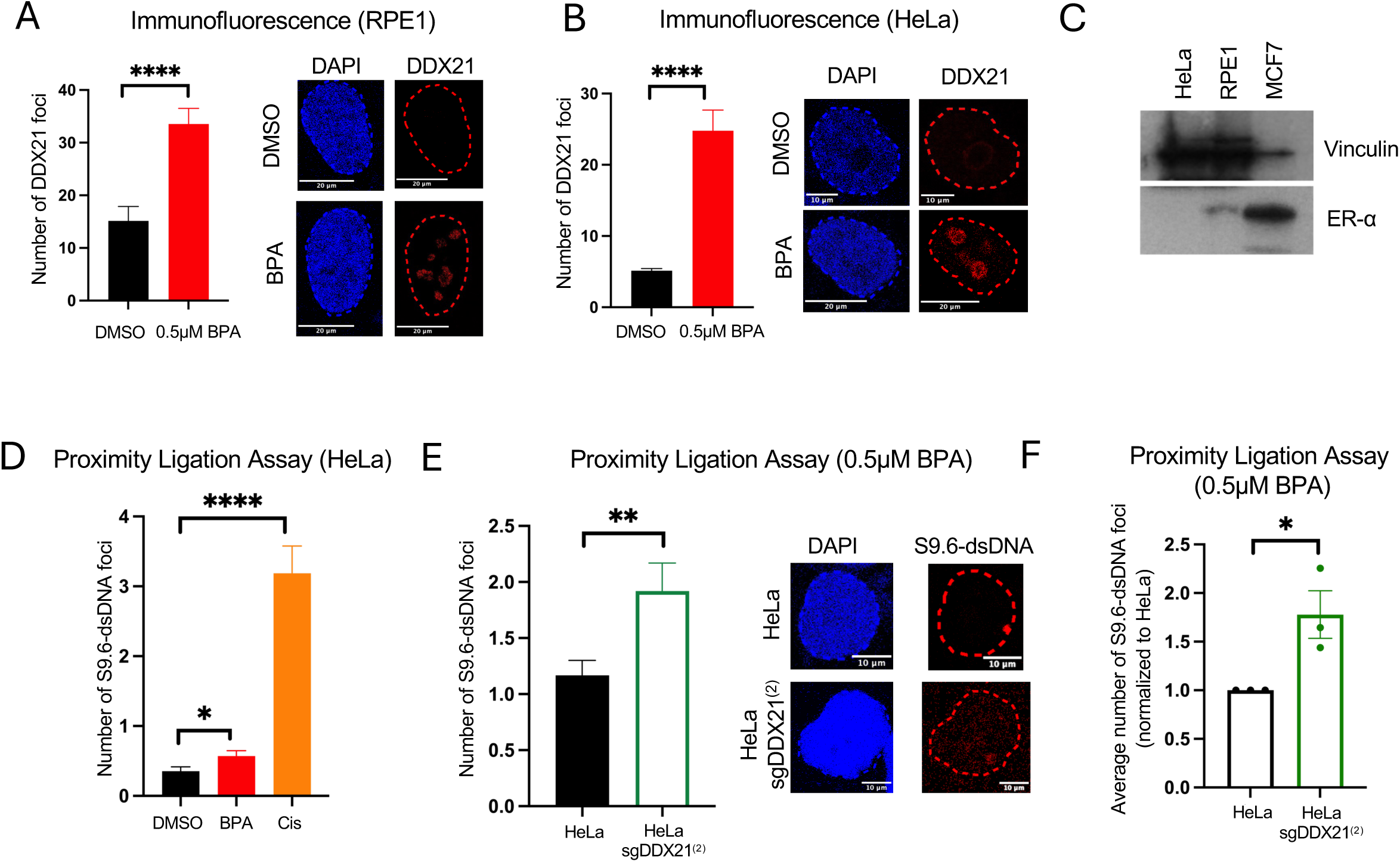
DDX21 suppresses BPA-induced R-loops in Estrogen receptor-negative cells. (**A, B**) DDX21 immunofluorescence showing that treatment with 0.5 μM BPA for 72 hours increases DDX21 foci formation in in RPE1 (**A**) and HeLa (**B**) cells. At least 105 cells were quantified for each condition. The mean value is represented on the graphs, and asterisks indicate statistical significance (t-test two-tailed, unpaired). Representative micrographs, with scale bars representing 10 and 20 µm are shown. (**C**) Western blot showing that HeLa and RPE1 cells do not express the estrogen receptor (ER). ER-expressing MCF7 cells used as a positive control. (**D**) Proximity ligation assay showing that treatment with 0.5 μM BPA for 72 hours or 0.5 μM Cisplatin for 24 hours increases the S9.6-dsDNA co-localization in HeLa cells. At least 101 cells were quantified for each condition. The mean value is represented on the graphs, and asterisks indicate statistical significance (t-test two-tailed, unpaired). (**E**) Representative PLA experiment showing an increase in the number of S9.6-dsDNA foci in sgDDX21 cells, as compared to control HeLa cells, after 0.5 μM BPA for 72 hours. The mean values are represented on the graphs, and asterisks indicate statistical significance (t-test two-tailed, unpaired). Representative micrographs, with scale bars representing 10 µm are shown. (**F**) The means of 3 independent S9.6-dsDNA PLA experiments showing an increase in the mean number of foci in sgDDX21 cells, normalized to control HeLa cells, after 0.5 μM BPA for 72 hours across three independent replicates. The mean values are represented on the graphs, and asterisks indicate statistical significance (t-test two-tailed, unpaired).

We thus next investigated if this DDX21 recruitment to DNA upon BPA exposure is connected to its R-loop suppression function. To this end, we first tested if BPA exposure increases R-loop formation in HeLa and RPE1 cells. BPA-induced R-loop formation was previously shown in estrogen receptor-expressing MCF7 cells upon treatment with 1-100ng/ml BPA and assigned to estrogen receptor-mediated transcriptional activation [10]. In contrast to MCF7, HeLa and RPE1 cells express no or very little estrogen receptor (Figure 6C). To measure R-loops, we employed the S9.6 antibody recognizing DNA-RNA hybrids. Since in our hands this antibody proved difficult to use in immunofluorescence studies, we instead measured R-loops by employing proximity ligation assays (PLA) with S9.6 and an antibody recognizing double stranded DNA (dsDNA). We found that treatment of HeLa cells with 0.5μM BPA for 3 days resulted in a small, but significant increase in the S9.6-dsDNA PLA signal (Figure 6D), arguing that exposure to low-dose BPA has the potential to increase the formation of R-loops in an estrogen receptor-independent manner. As control, treatment with the DNA damaging agent cisplatin shows a strong increase in R-loop formation, as previously described [28].

We next investigated the impact of DDX21 depletion on BPA-induced R-loop formation. PLA assays showed that sgDDX21 HeLa cells have increased R-loop formation upon treatment with 0.5μM BPA for 3 days (Figure 6E). This result was confirmed in 3 independent experiments (Figure 6F). Overall, these findings show that environmentally relevant exposure to low-dose BPA could cause an estrogen receptor-independent increase in R-loops. The RNA helicase DDX21 is critical to suppress these R-loops, potentially explaining why loss of DDX21 sensitizes cells to BPA under these conditions.

## Discussion

While BPA exposure was associated with numerous adverse health effects, including abnormalities in reproductive organ function, placental dysfunction, precocious puberty, and neurological impairment, its association with carcinogenesis is less clear. While a number of studies indicated that BPA exposure increases proliferation, those studies were generally performed at much higher BPA concentrations compared to the amount of BPA found in the serum or urine of the general population. Mechanistically, BPA exposure was shown to increase the transcription of oncogenes, as well as cause R-loops and DSBs. However, many of these effects were assigned to activation of estrogen signaling. Independent of ER signaling, BPA exposure has been shown to cause DNA adducts. However, as mentioned above, these studies were performed at BPA concentrations far exceeding the BPA dose found in people, including those with increased exposure to BPA such as plastics manufacturing workers [7].

In our study, we aimed at identifying genetic mechanisms responding to physiologically relevant BPA exposure, and in particular investigating if DNA repair is one of them, through an unbiased whole-genome CRISPR screening approach. A recent study [29] reported a BPA sensitivity CRISPR screen, however that screen was performed with a BPA concentration of 75μM, which is about 7,500-fold higher than the mean BPA concentration in the urine or serum of the general population. In contrast, we performed BPA sensitivity CRISPR screens using a concentration of BPA similar to that found in the serum and urine of plastics manufacturing workers, namely 0.5μM. For increased rigor, we performed our screen in two different cell lines, namely cervical cancer HeLa cells and non-transformed epithelial RPE1 cells. Surprisingly, there was a somewhat limited overlap between the results in the two cell lines, with only 64 genes found in common among the top hits with MAGeCK score lower than 0.01, compared to the expected random probability of overlap between the two datasets, which is 28.

Nevertheless, under these conditions, we found that biological pathways linked to DNA homeostasis were the only ones common between the two cell lines (including DNA repair, cell cycle, sister chromatid cohesion). This suggests that exposures to physiologically-relevant BPA concentration may cause DNA damage. Indeed, we validated the double strand break repair gene RAD51C and the R-loop resolvase DDX21 as BPA sensitivity suppressors. We moreover validated two other top hits common between the HeLa and U2OS screens, namely ATG9A and CPSF2.

Beyond validating RAD51C as a BPA sensitivity gene, our study shows that BPA exposure at this relevant concentration can cause DNA damage and/or replication stress, which we and others previously showed using higher BPA concentrations [13,14]. RAD51C is inactivated in Fanconi Anemia patients, as well as patients with familial breast and ovarian cancer predisposition syndromes [30]. Our study thus suggests that individuals with mutations in homologous recombination genes may have increased sensitivity to the detrimental effects of BPA exposure. Moreover, while we observed a reduction in γH2AX foci formation in RAD51C knock-down cells exposed to BPA as compared to DMSO, our experiments could not distinguish whether this was due to an alteration in DNA damage formation or a defect in DNA damage signaling. Future directions include assessing ATM/ATR/CHK2 pathway activation in the presence and absence of RAD51C in response to BPA exposure.

Our study also sheds additional light on how BPA exposure may cause DSBs. Previous studies indicated that BPA treatment causes DNA adducts [15,16]. These adducts can lead to replication fork stalling and collapse, generating DSBs. Our laboratory previously showed that BPA treatment also causes ssDNA gaps, presumable upon the restart of replication forks arrested at BPA adducts by the primase polymerase PrimPol downstream of the adducts [14]. These studies were performed at high BPA concentration, and thus the extent to which these events occur upon exposure to physiologically relevant BPA doses remains unclear. Here, we show that low-dose BPA exposure can result in the formation of R-loops, which are known to eventually cause DSB formation unless timely resolved.

Previous studies showed that BPA exposure causes the formation of R-loops in ER-positive cells, presumably through activation of ER-mediated transcription. However, we show here that low-dose BPA treatment can potentially result in R-loop formation in ER-negative cells as well. These R-loops may thus be unlikely to form from increased transcriptional activity, but instead they may form at sites of BPA adduct formation, as previously shown for alkylating adducts [31]. Regardless of their source, it seems likely that R-loop formation by BPA would be amplified in ER-positive cells such as the breast tissue due to concurrent ER activation. Interestingly, other environmental genotoxic agents, such as Benzu[a]pyrene, were previously shown to cause R-loops[32], suggesting that R-loop repair may be critical to suppressing genomic instability induced by environmental agents. Limitations of our study include complimentary validation of the R-loop formation observed in Figure 6. Future directions include assessing how the presence or absence of DDX21 affects transcription–replication conflict formation after BPA exposure.

In addition, we also identified ATG9A and CPSF2 as potential BPA sensitivity genes. ATG9A is an autophagy protein involved in phagophore formation and lipid mobilization [33]. Since these functions are associated with cellular membrane protection, it is possible that loss of ATG9A improves the cellular intake of BPA. CPSF2 is a critical factor involved in mRNA polyadenylation, and regulates alternative splicing [34,35]. Thus, loss of CPSF2 could indirectly affect BPA sensitivity by changing the composition of the proteome. Moreover, loss of CPSF2 expression has been associated with worse clinical outcome in patients with papillary thyroid carcinoma[36,37]. Given that BPA is a genotoxic agent, understanding how loss of CPSF2 promotes cellular sensitivity to BPA may also help in the understanding of treatment response in papillary thyroid carcinoma.

Even though the concentration used is physiologically relevant, our study is likely to under-represent the toxic effects of BPA considering that we only treated cells for 19 days for the BPA sensitivity screens. It is likely that the effects we glimpsed in our study are amplified multi-fold in plastics manufacturing workers exposed to BPA over many years. Moreover, steady state concentrations in plastic manufacturing workers were found to range all the way to 144μM [3]. In addition, studies showed that levels of BPA in sweat are detectable even when no detectable BPA levels were found in urine or serum [4], suggesting bioaccumulation that is not accounted for in reports of physiologically relevant BPA concentrations (which measure BPA levels in urine and/or serum). Overall, these findings highlight the need to devise better studies in order to improve the occupational health safety guidelines for these individuals, as well as regulate the BPA contents in every-day plastic items to limit the BPA exposure of the general population.

## Acknowledgments

We would like to thank Dr. Alan D’Andrea, Dr. Ashna Dhoonmoon, Dr. Jude Khatib, and Justyn Dusold for materials and support, and the Penn State College of Medicine Advanced Light Microscopy (RRID:SCR_022526), Genome Sciences (RRID:SCR_021123) and Flow Cytometry (RRID:SCR_021134) core facilities.

## Legends to Supplementary Tables

**Supplementary Table S1.** MAGeCK analyses of the CRISPR screens identifying genes whose loss confers increased sensitivity to BPA in HeLa cells.

**Supplementary Table S2.** MAGeCK analyses of the CRISPR screens identifying genes whose loss confers increased sensitivity to BPA in RPE1 cells.

**Supplementary Table S3.** List of common hits with MAGeCK score lower than 0.01 from the CRISPR knockout screens for BPA sensitivity in HeLa and RPE1 cells.

